# Discovery of broadly-neutralizing antibodies against brown recluse spider and Gadim scorpion sphingomyelinases using consensus toxins as antigens

**DOI:** 10.1101/2023.04.17.537284

**Authors:** Esperanza Rivera-de-Torre, Stefanos Lamparidou, Markus F. Bohn, Seyed Mahdi Kazemi, Andreas H. Laustsen

## Abstract

Broadly-neutralizing monoclonal antibodies are becoming increasingly important tools for treating infectious diseases and animal envenomings. However, designing and developing broadly-neutralizing antibodies can be cumbersome using traditional low-throughput iterative protein engineering methods. Here, we present a new high-throughput approach for the standardized discovery of broadly-neutralizing monoclonal antibodies relying on phage display technology and consensus antigens representing an average sequence of related proteins. We showcase the utility of this approach by applying it to toxic sphingomyelinases from the venoms of very distant orders of the animal kingdom, the recluse spider and Gadim scorpion. First, we designed a consensus sphingomyelinase and performed three rounds of phage display selection, followed by DELFIA-based screening and ranking, and benchmarked this to a similar campaign involving cross-panning against recombinant versions of the native toxins. Second, we identified two scFvs that not only bind the consensus toxins, but which can also neutralize sphingomyelinase activity *in vitro*. Finally, we conclude that the phage display campaign involving the use of the consensus toxin was more successful in yielding cross-neutralizing scFvs than the phage display campaign involving cross-panning.

## Highlights

- Broadly-neutralizing antibodies are useful agents for targeting multiple antigens
- Consensus toxins represent an average sequence of multiple native toxins
- Broadly-neutralizing antibodies can be discovered using consensus toxins and *in vitro* selection
- Consensus sphingomyelinases were designed and used for discovery of broadly-neutralizing antibodies

## Introduction

Monoclonal antibodies (mAbs) are the largest group of biotherapeutic agents used to treat diseases, with more than 100 approved for clinical applications ^1^. However, for some applications, such as controlling infectious diseases or neutralizing toxins, the high specificity of mAbs comes with the disadvantage that mAbs may not always be robust against antigenic variation ^2^. This limitation has recently been a particular concern for mAbs targeting escape mutants of SARS-CoV2 ^3^, but antigen variation has for more than a century also complicated the development of envenoming therapies, where antibodies need to target large panels of similar and dissimilar toxins ^4^. Human mAbs owe their success to their compatibility with the human immune system and their versatility, but broadening their neutralization capacity beyond a single epitope or even a single protein may be critical for the treatment of rapidly evolving or multi-component diseases.

Recently, it has been demonstrated that broadly-neutralizing mAbs (bnAbs) targeting animal toxins can be developed using cross-panning strategies in phage display selection campaigns ^5, 6^, by semi-rational design and directed evolution ^7^, and by high-throughput screening of B-cells from immunized individuals ^8^. Here, we demonstrate the utility of a fourth strategy that uses consensus toxins, which are artificial toxins based on the average sequence of related toxins^9^. These have previously been used as immunogens to generate broadly-neutralizing polyclonal antibody responses in animals ^10, 11^. The objective of this study was to explore the utility of consensus antigens in an *in vitro* display technology-based antibody discovery campaign ^12, 13^. This was achieved by designing a consensus toxin and using phage display selection to discover bnAbs with cross-reactivity against toxins from two distantly related species deriving from distant orders of the animal Kingdom, namely *Loxosceles sp.* (recluse spider) and *Hemiscorpius sp.* (Gadim scorpion) found in North Africa and the Middle East, respectively ^14^, and comparing this discovery campaign to one utilizing a previous established cross-panning strategy ^5^.

Both recluse spiders and Gadim scorpions are of medical importance due to the high incidence of scorpion stings in the region. In Iran alone, there are between 40,000 and 50,000 cases of scorpion stings annually, resulting in approximately 20 deaths, with *H. lepturus* responsible for 10% to 25% of all sting cases and nearly 70% of the fatalities ^15^. Unlike other scorpion stings, *Hemiscorpius sp*. stings are not immediately painful, but result in delayed necrotoxic and hemotoxic symptoms that resemble those caused by *Loxosceles sp*. bites. Due to the delay in the onset of clinical manifestations, victims may delay seeking medical attention, which in turn contributes to the high mortality rate associated with *Hemiscorpius sp*. stings. The primary toxin responsible for the clinical manifestations observed for *Hemiscorpius sp.* stings is a sphingomyelinase (SMase), for which related isoforms also exists in *Loxosceles sp*. venom. Both SMases cause cytotoxicity by transforming sphingomyelin in cellular membranes into ceramide and phosphorylcholine, thereby disrupting the membranes and triggering necrosis through ceramide-induced apoptosis ^16–18^. Therefore, neutralizing these SMases early upon envenoming is of high relevance to achieve optimal treatment outcomes for bite victims.

In this work, we designed a consensus SMase (cSMase) that is an artificial toxin resembling an average sequence and structure of both *Loxosceles sp.* and *Hemiscorpius sp.* SMases. We demonstrated the advantage of using cSMase as an antigen in a phage display selection campaign, discovering single-chain variable fragments (scFvs) that bind and neutralize native SMases from both species. We compared the utility of a stringent selection strategy using cSMase as an antigen with a cross-panning strategy using recombinant versions of the native toxins and found that the strategy involving the consensus toxin was more successful as it yielded scFvs that bind to natural SMases from *H. lepturus* and *L. rufescens* with higher affinity than those scFvs identified using the cross-panning strategy. Furthermore, we demonstrate that the scFvs obtained using cSMase bind to SMases from *H. lepturus* and *L. rufescens* with similar affinity and can effectively neutralize the Smase activity of *H. lepturus* whole venom. These results provide evidence for the potential utility of using consensus antigens and phage display technology for discovery of broadly-neutralizing antibodies.

## Results

For the design of a consensus sphingomyelinase (cSMase), a blastp search was performed on the NCBI database using the most representative queries from scorpions and recluse spiders. This search resulted in 136 unique sequences based on non-duplication and coverage criteria (>70%) (File S1). These sequences were aligned in Jalview 2, and a cSMase was designed based on the most commonly present amino acid or physicochemical property on each position (acid, basic, polar, or hydrophobic amino acid side chains). The signal peptides were manually trimmed from the final sequence to obtain the mature protein. Although designed cSMase was not identical to any of the natural sequences, it shared 47.60% and 86.06% similarity with the most representative SMases from *H. lepturus* (A0A1L4BJ98) and *L. rufescens* (C0JB02.1), respectively (Fig. 1A), which are the most prevalent Gadim scorpions and recluse spiders in the Middle East and North Africa region. Despite the differences in primary sequence, AlphaFold2 predicated that the three-dimensional structures of the two natural SMases and the designed cSMase would be similar, featuring a β-sheet barrel surrounded by α-helices (Fig. 1B).

**Figure 1.**
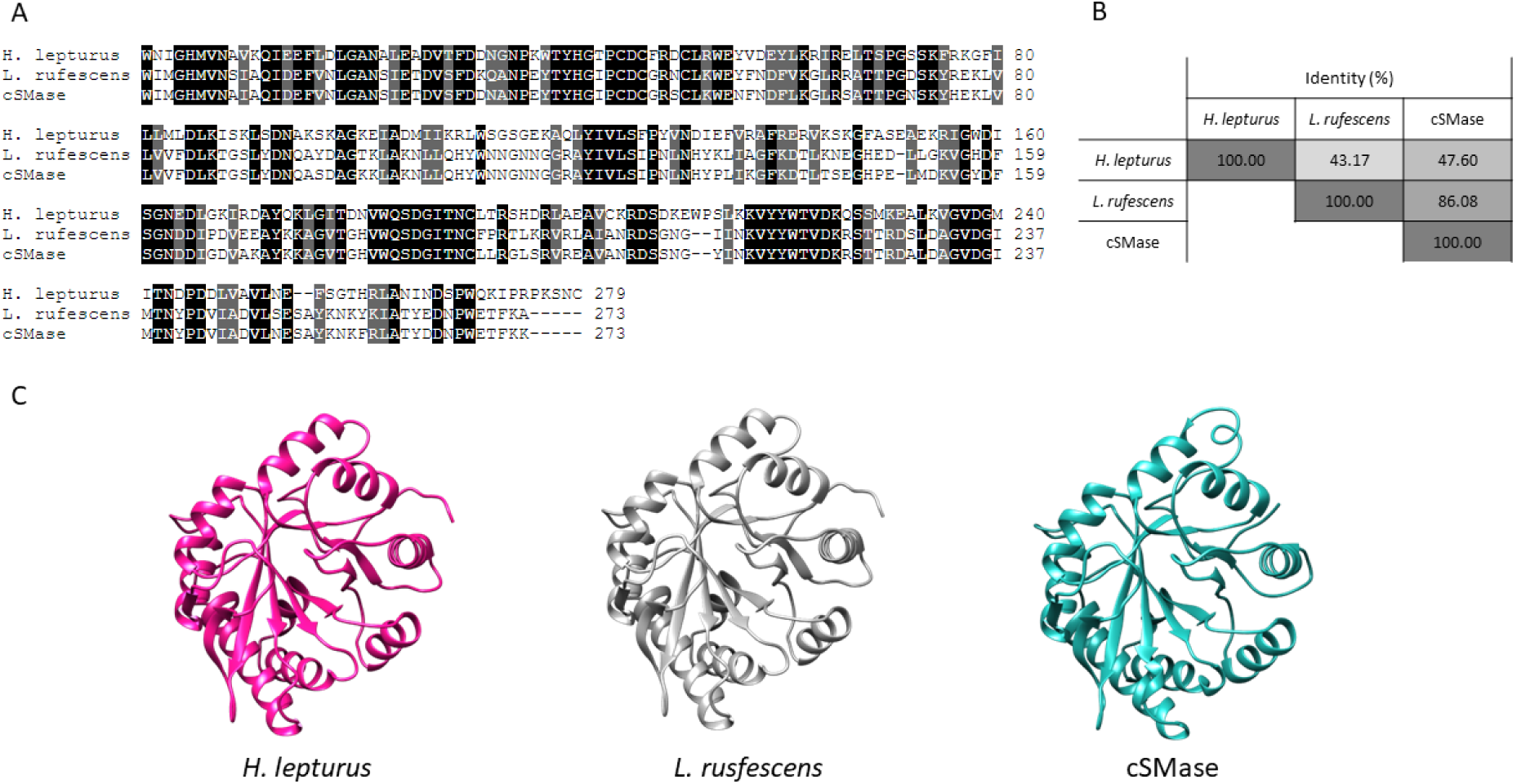
Comparison of SMases from H. lepturus (A0A1L4BJ98), L. rufescens (C0JB02.1), and the designed consensus SMase (cSMase). (A) Sequence alignment was performed using Clustal Omega default settings, and the resulting alignment was color-coded to indicate the degree of conservation for each position. A black shade indicates a completely conserved amino acid, while a grey shade indicates conservation of physicochemical properties. White indicates non-conserved amino acids. (B) The percentage identity matrix shows the level of similarity between the aligned proteins. The values are indicated as percentages and are displayed in a color-coded matrix. (C) The predicted three-dimensional structure of cSMase was generated using AlphaFold2.

The coding sequences of the *H. lepturus* (A0A1L4BJ98), *L. rufescens* (C0JB02.1), and the cSMase were optimized for bacterial codon-usage and subcloned into an expression vector in phase with the 6xHis tag. The recombinant proteins were expressed in BL21(DE3) *E. coli* and purified to homogeneity with a single affinity chromatography step. *H. lepturus* venom was fractionated via HPLC and fractions exhibiting electrophoretic mobility on SDS-PAGE consistent with SMases were selected for further analysis. Circular dichroism spectra of the recombinant toxins and the SMase purified from *H. lepturus* venom showed that the pure proteins had a folded structure with a relative minimum ellipticity at 220 nm and an absolute minimum value around 208 nm, which is a hallmark of α-helix-rich structures (Figure 2). These spectra closely resembled those of native SMases purified from the crude venom of *Loxosceles sp.* ^19^. Thermal stability of the recombinant proteins was tested using a differential scanning fluorometer (NanoDSF) and the T_m_ values were determined by fitting the experimental data to a polynomial function. The slope maxima indicated by the peak of its first derivative resulted in 72.3 °C, 71.63 °C, and 72.6 °C for rLr, rHl, and cSMase respectively. All the proteins were able to refold, at least partially, upon cooling with T_m_ values equivalent to the first ones measured for all the proteins. The T_m_ for the SMase purified from *H. lepturus* crude venom (Hl) was determined to be 72.6 °C, and the protein was also able to refold upon cooling. The differences between the Hl and the recombinant proteins might be due to minor contaminations from other components of the venom in the fraction or post-translation modifications that are absent in the recombinant proteins, which could influence in the thermal stability.

**Figure 2.**
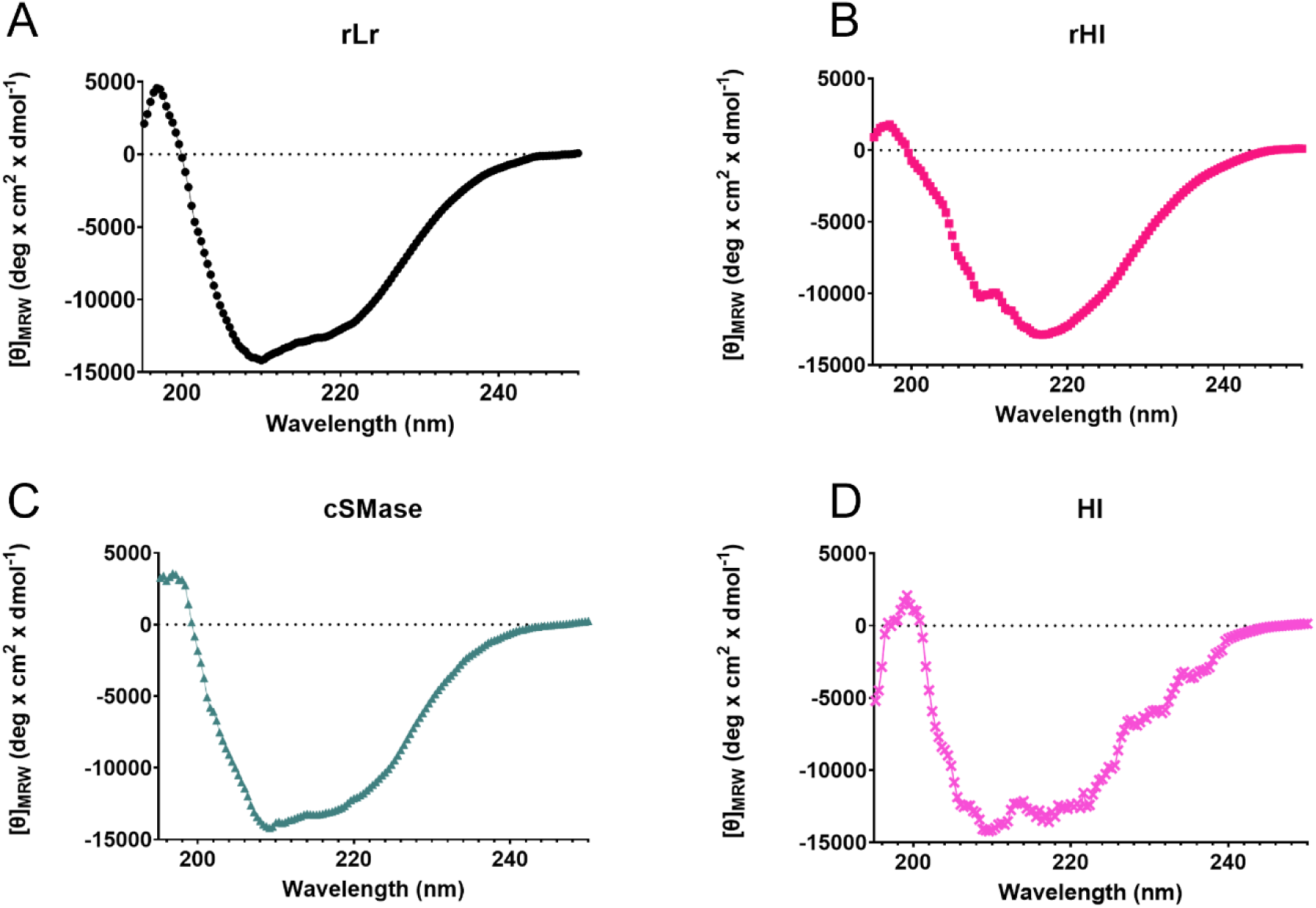
Far-UV circular dichroism spectra of rLr (A - black dots), rHl (B - magenta squares), cSMase (C - green triangles), and Hl purified from H. lepturus venom (D - pink crosses).

Phage display technology was employed to select scFv antibody fragments from the IONTAS phage library κ with the aim of obtaining broadly-neutralising antibodies against scorpion and spider toxins. Two strategies were used to achieve this: cross-panning with recombinant toxins (rLr and rHl) and stringent panning with the cSMase. To enrich the pool of phages cross-binding to rLr and rHl, three rounds of cross-panning were performed. In the first round, biotinylated rLr or rHl was used as antigen at concentrations of 100 nM. The antigen was alternated in the second round of panning but kept at the same concentration of 100 nM. However, no enrichment of phages binding to the recombinant toxins was observed in the second round, indicating a lack of cross-binding of the antibody fragments displayed in the phage pools (Fig 4A and 4B). In the third round of selection, the concentration of the antigen used in the second round was reduced to 50 nM, resulting in a slight enrichment of phages binding to the target used in the first round. Specifically, compared to the second round, the third round showed 2.5x and 2.75x enrichment for the selections starting with rLr and rHl, respectively.

**Figure 3.**
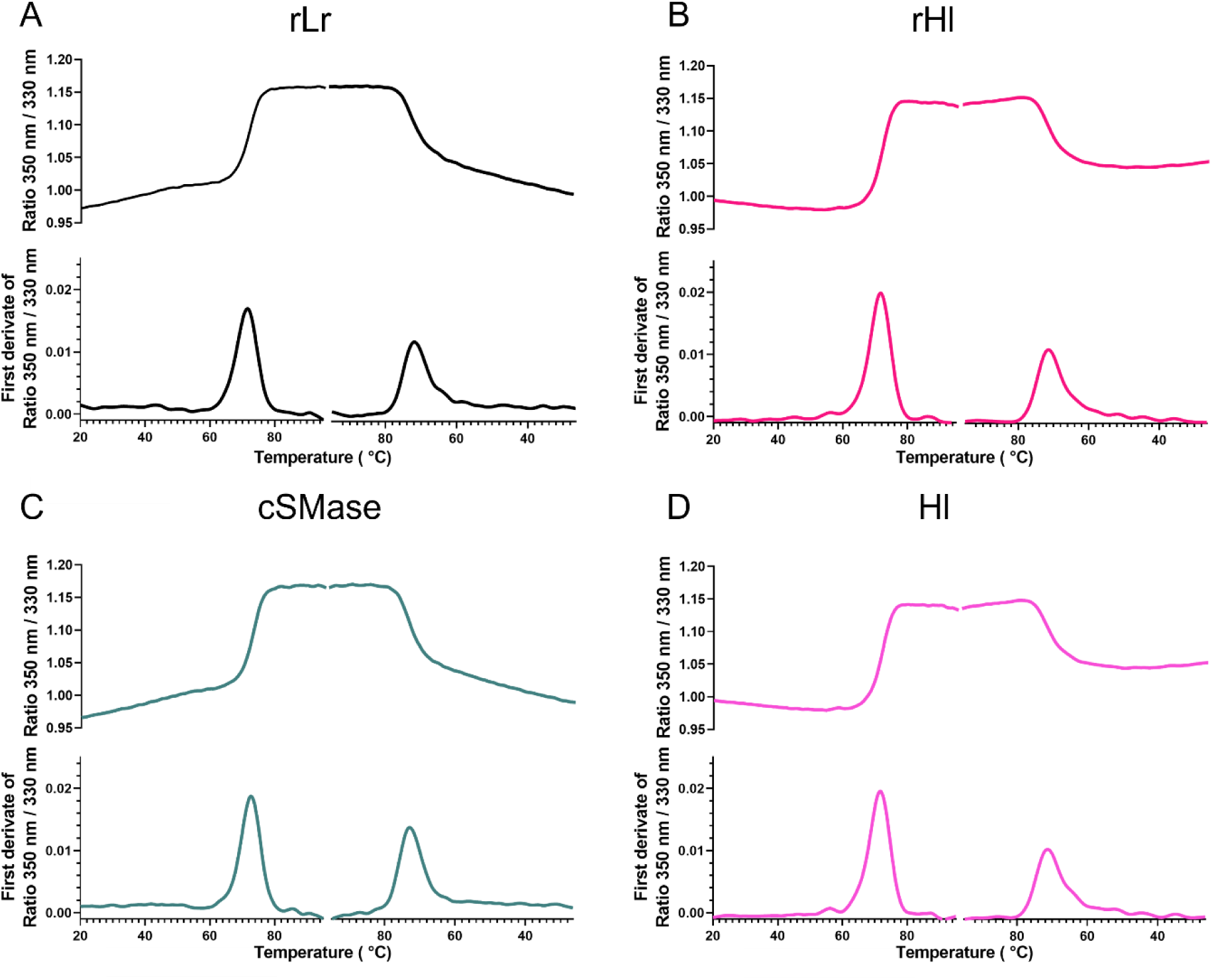
Thermal denaturation curves in NanoTemper for rLr (A - black), rHl (B - magenta), cSMase (C - green), and Hl (D - pink).

**Figure 4.**
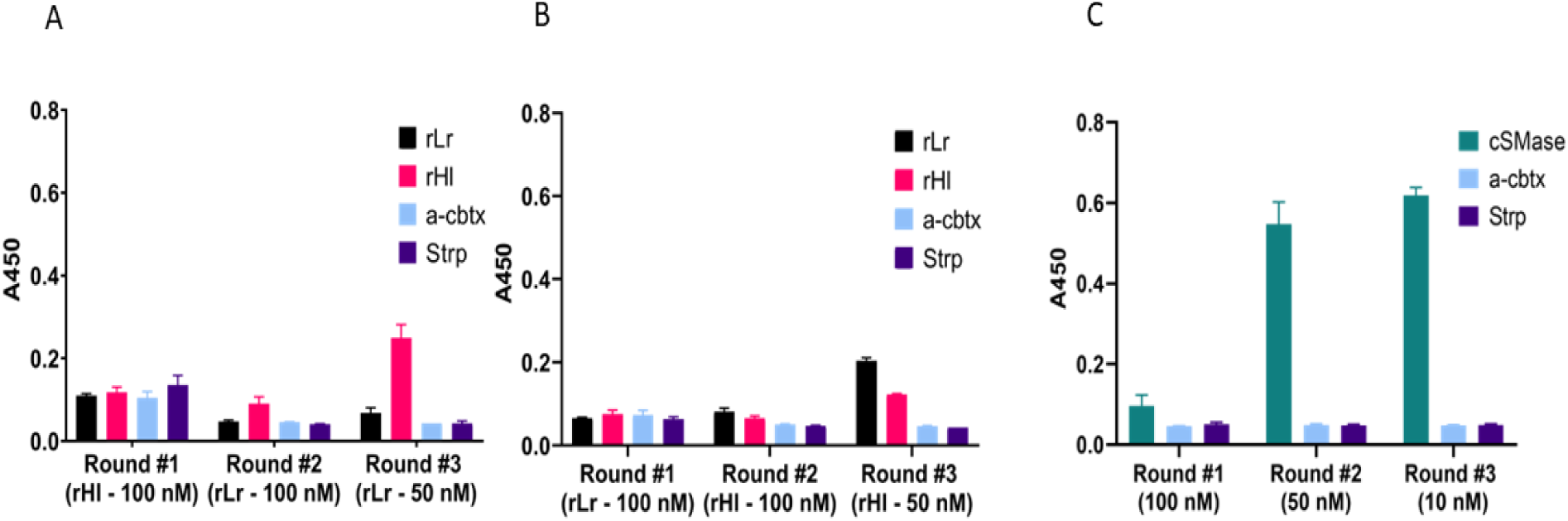
Polyclonal phage ELISA of the outputs from each phage display round (A, B, C). (A) Cross-panning with recombinant toxins (rLr 100 nM, rHl 100 nM, rHl 50 nM). (B) Cross-panning with recombinant toxins (rHl 100 nM, rLr 100 nM, rLr 50 nM). Stringent panning with cSMase (100 nM, 50 nM, 10 nM).

The second strategy involved the use of cSMase as the antigen for all three rounds of panning, with decreasing antigen concentrations in each round (100 nM, 50 nM, and 10 nM). This approach aimed to select for high-affinity cross-binders. The strategy resulted in an enrichment of phages displaying scFvs specifically binding to the cSMase in the second and the third round (Figure 4C).The scFv-encoding genes in the phages from the third panning round performed with cSMase were isolated, sub-cloned into the pSANG10-3F expression vector ^20^, and 276 clones were picked.

The recombinant monoclonal scFvs were expressed in solution and tested for their ability to bind biotinylated rLr or rHl using a Time-Resolved Fluorescence (TRF) assay. The TRF signal, which is directly proportional to the binding strength of each scFv clone, was recorded and analyzed. The scatterplot in Figure 5 displays the TRF signals for the ability of each scFv clone to bind to both recombinant toxins, rHl and rLr. It is worth noting, that some scFvs showed equal binding intensities to both rHl and rLr. To identify the most optimal cross-binding scFvs, a cut-off value of 3 x 10^4^ TRF intensity over the negative control for both rHl and rLr and a TRF rLr/TRF rHl ratio between 0.8 and 1.2 was used (Table 1). As a result, two scFv binders were chosen for further characterization through DNA sequencing, resulting in two unique sequences. The scFv binders TPL0674_03_A04 and TPL0674_03_F05 were produced in a mg scale and purified for further investigation of their binding affinity and neutralisation capacity.

**Figure 5.**
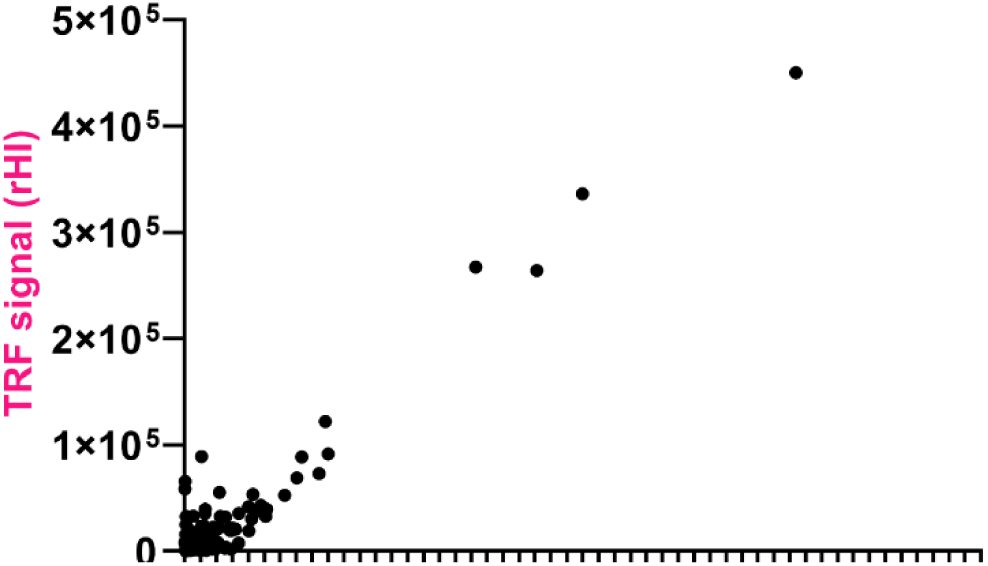
Monoclonal DELFIA. Binding to rLr (x axis) and rHl (y axis). Each dot represents a single monoclonal scFv.

**Table 1.**
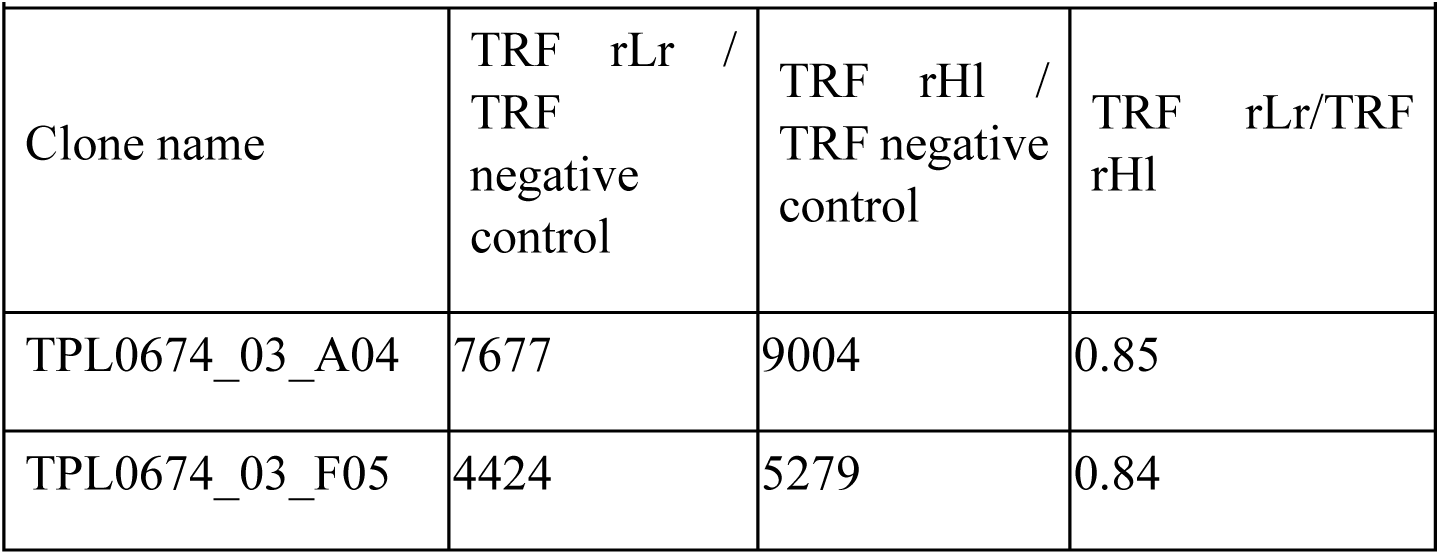
Binders selected for sequencing

| Clone name | TRF rLr /<br>TRF<br>negative<br>control | TRF rHI /<br>TRF negative<br>control | TRF rLr/TRF<br>rHI |
| --- | --- | --- | --- |
| TPL0674_03_A04 | 7677 | 9004 | 0.85 |
| TPL0674_03_F05 | 4424 | 5279 | 0.84 |

The binding affinity of cross-reactive scFv antibodies, including recombinant SMases (rLr and rHl) and a biotinylated natural SMase purified from *H. lepturus*, was assessed using biolayer interferometry. The results demonstrated comparable binding affinity in the nanomolar range for both scFvs against the recombinant and natural toxins (Figure 6 and Table 2).

**Figure 6.**
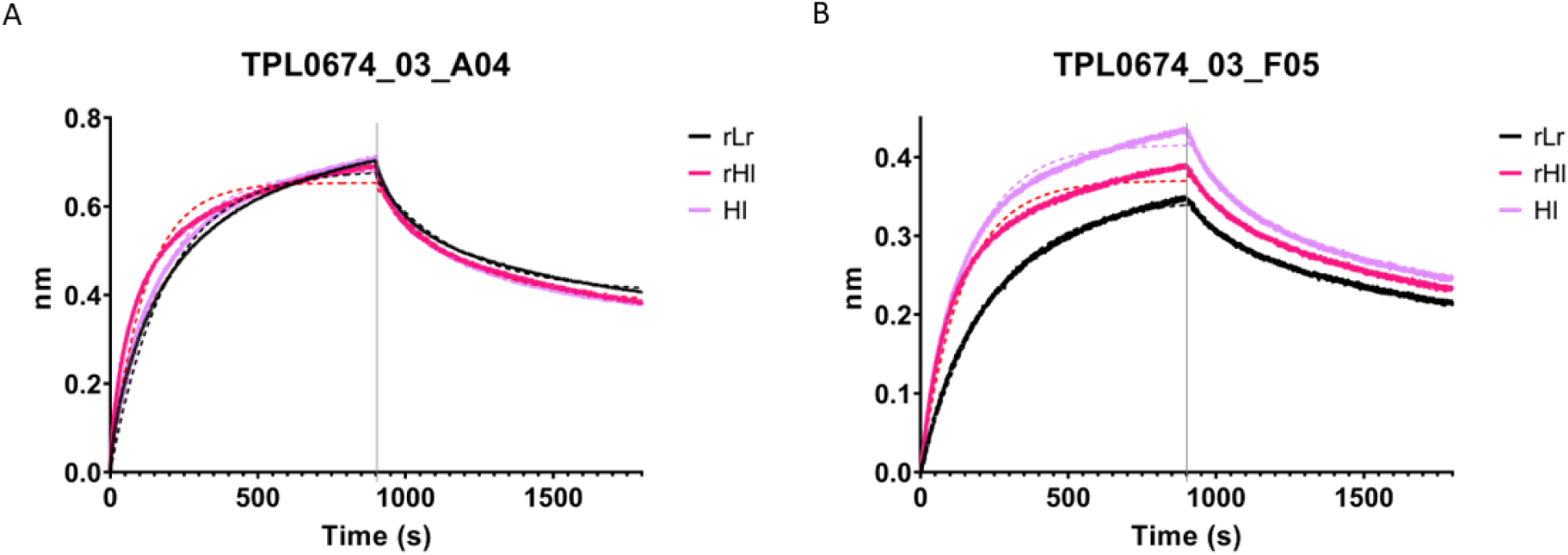
Binding curves measured by biolayer interferometry of TPL0674_03_A04 clone (A) and TPL0674_03_F05 clone (B) to rLr (black), rHl (magenta), and Hl purified from crude venom (pink).

**Figure 7.**
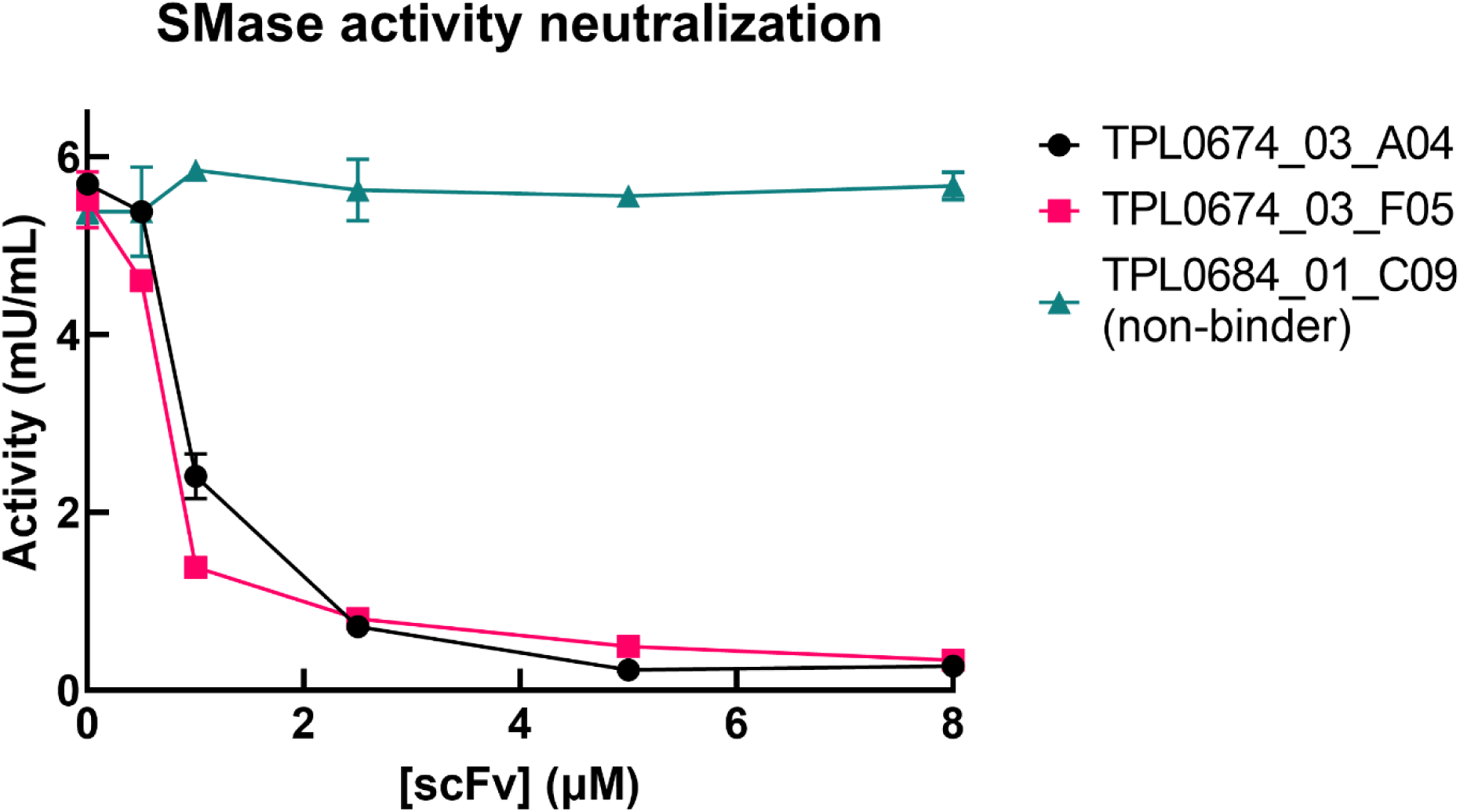
Neutralization of SMase activity of H. lepturus whole venom by TPL0674_03_A04 (black dots) and TPL0674_03_F05 (pink squares). An unspecific scFv (green triangles) was used as control.

**Table 2.** Binding affinity determined by biolayer interferometry

| Table 2. Binding affinity determined by biolayer interferometry |  |  |
| --- | --- | --- |
| Clone | Toxin | $K_D \pm SD$ (nM) |
| TPL0674_03_A04 | HI | $70.3 \pm 0.9$ |
| | rLr | $66.9 \pm 0.8$ |
| | rHI | $45.3 \pm 0.6$ |
| TPL0674_03_F05 | HI | $60.7 \pm 0.6$ |
| | rLr | $80.4 \pm 0.7$ |
| | rHI | $51.6 \pm 0.5$ |

To determine whether the cross-reactive scFvs could neutralize toxicity, the enzymatic SMase activity of *H. lepturus* venom was tested in the presence and absence of the scFvs, at concentrations ranging from 0.5-8 µM. An irrelevant scFv (TPL0684_01_C09) was used as negative control. Both scFvs (TPL0674_03_A04 and TPL0674_03_F05) were found to successfully inhibit the SMase activity of *H. lepturus* venom, while the irrelevant scFv did not have any effect on the SMase

In summary, the results demonstrated that the designed consensus sphingomyelinase (cSMase) could be successfully used to select cross-reactive scFv antibodies (TPL0674_03_A04 and TPL0674_03_F05) that bind to recombinant and natural toxins with comparable affinity. These scFvs were found to specifically inhibit the SMase activity of *H. lepturus* venom, highlighting their potential as antibody leads for the development of broad-spectrum toxin neutralizing agents. Based on these observations, we demonstrate that consensus proteins can be used as antigens during phage display selection campaigns for the selection of cross-binding/neutralizing antibodies. While these results showcase the utility of this approach on SMases from scorpion and spider venoms, we speculate that this methodology might be applicable for other targets for the development of therapeutic antibodies.

## Discussion

In this study, we present the identification of several human monoclonal scFvs with broad neutralization activity, obtained using phage display technology. The scFvs were selected against a recombinant toxin that was designed to mimic a consensus of the SMases produced by *Hemiscorpius sp.* and *Loxosceles sp.* The identified scFvs could bind to recombinantly produced version of native SMases from *H. lepturus* and *L. rufescens*, as well as a natural SMase purified from *H. lepturus* venom, with comparable affinities. Furthermore, the cross-binding scFvs could neutralize the SMase activity of *H. lepturus* whole venom *in vitro*. We found that the cross-panning strategy was not effective in generating broadly-neutralizing scFvs for the target toxins we examined, as there was a lack of cross-reactivity for the phage outputs. This suggests that the target toxins might not share enough epitope similarity to enable the obtainment of cross-binders in a cross-panning campaign, as rHl and rLr only share 43% sequence identity. However, we did find that the use of the cSMase, which represents the average of more than 100 native SMases, consistently enabled the enrichment of binding phages in each round of selection under stringent conditions, *i.e.*, increasing selection pressure through successive decreases in antigen concentration during consecutive panning rounds.

Consensus toxins have previously been used to immunize rabbits and a horse and isolate serum containing polyclonal mixtures of antibodies displaying broad-spectrum neutralization capabilities ^10, 11^. However, it remains unclear if these broad neutralization capabilities derived from the induction of monoclonal bnAbs in the polyclonal mixture or if the broad neutralization capabilities derived from the polyclonality of the antiserum^21^. Nevertheless, based on the data presented here, we can assert that consensus toxins can serve as powerful tools for obtaining monoclonal scFvs that can neutralize quite dissimilar antigens from distant sources.

The clinical relevance of envenoming therapies against scorpion stings and spider bites is particularly significant in resource-poor areas of the world ^14^. Diagnosing the specific spider or scorpion responsible for a bite can be challenging, underscoring the need for broadly effective envenoming therapies. Our study shows that the same scFv can neutralize similar toxins from very distantly related species, which demonstrates its potential utility as a broadly-neutralizing therapeutic agent for arachnid envenomings, if the results observed *in vitro* translate into the *in vivo* setting.

The use of consensus antigens, as presented here, can likely be extended to other areas, where the development of broadly-neutralizing antibodies could be of therapeutic relevance. These areas include envenomings by other venomous animals, such as snakes or cone snails, infectious diseases, such as influenza or HIV, or hypermutating disease targets, such as those found in many cancers ^22–24^. Moreover, we envision that the use of consensus antigens may also find applications in combination with other established methods for the discovery and design of broadly-neutralizing antibodies, including cross-panning-based phage display campaigns, structural approaches, and hybridoma and B-cell screening, and thereby expand the toolbox for the antibody discoverer.

## Materials and methods

### Consensus toxin design

Amino acid sequences for SMases from Hemiscorpiidae (scorpions) and Sicariidae (recluse spiders) families were collected from the National Center for Biotechnology Information (NCBI) website via blastp using as query the most representative members of the toxin family (A0A1L4BJ98 from *H. lepturus*, P0CE80 from *L. Intermedia*, P0CE79.1 from *L. reclusa,* and C0JB02.1 from *L. rufescens*) and default local alignment parameters. All hits were pooled and curated to eliminate duplicated entries. Multiple sequence alignments of the SMases were performed using the Jalview 2 program, with the signal peptides trimmed manually from all the sequences ^25^. Sequences that overlapped < 70% of the sequence length were eliminated from the pool. Consensus sequences were determined by conserving amino acid residues or physicochemical properties for each position.

### Molecular modeling of cSMase structure

The three-dimensional (3D) structures of cSMase and *L. rufescens* SMase were predicted using the state-of-the-art AlphaFold2_mmseqs2 algorithm, which is part of the ColabFold set of Jupyter notebooks. The detailed methodology for this approach can be found in the respective literature^26, 27^ Briefly, the amino acid sequence of cSMase was used as input for the AlphaFold2_mmseqs2 Jupyter notebook. This notebook is designed to predict protein structures using the AlphaFold2 algorithm, which combines machine learning techniques with multiple sequence alignment (MSA) generation using the mmseqs2 method. The algorithm employs deep learning techniques to identify patterns in the input data and produce a high-quality structural model. A predicted model of the *H. lepturus* SMase 3D structure was retrieved from the Alphafold Protein Structure Database^28^ and subjected to N-terminal truncation of the first 45 residues using phenix.pdbtools^29^.

### Cloning

The coding sequences for the native SMases from *H. lepturus* (A0A1L4BJ98) and *L. rufescens* (C0JB02.1), as well as the designed cSMase were codon-optimized for expression in *Escherichia coli* and purchased from Eurofins Genomics. All genetic constructs included the sequences *NcoI-* 6xHis and STOP-*BamHI* in the 5’ and 3’ ends. The genetic constructs were subcloned into the pET 6xHis TEV cloning vector (Addgene #29653) using standard restriction and ligation cloning procedures. Briefly, both plasmid and genetic constructs were digested with *NcoI* and *BamHI* restriction enzymes (FastDigest, Thermofisher), inactivated (5 min at 65 °C), and agarose gel purified (GeneJET Gel Extraction Kit). Purified fragments were mixed in a 1:5 ratio (vector:insert) for ligation with T4 ligase, following the manufacturer recommendation (New England Biolabs). *E. coli* 10G Chemically Competent Cells (Lucigen) were transformed with the ligation mixtures and plated on LB plates containing 50 µg kanamycin (LB-Kan) (Sigma-Aldrich). Plasmids from four colonies with the expected electrophoretic pattern after analytical digestion with *NcoI*/*BamHI* (FastDigest, Thermofisher) were Sanger-sequenced by Eurofins Genomics. The constructs were named pET-6xHis-rHl, pET-6xHis-rLr, and pET-6xHis-cSMase.

### Expression and purification of recombinant toxins

Constructs of the recombinant SMases with confirmed sequence via Sanger sequencing were transformed by heat-shock into *E. coli* BL21(DE3) competent cells for expression following the manufacturer’s instructions (New England Biolabs). Colonies harbouring pET-rHl, pET-rLr, and pET-cSMase were selected based on their ability to grow on LB-Kan plates (Sigma). Initial bacterial inoculum was grown overnight at 37 °C in baffled Erlenmeyer flasks in LB-Kan medium. Next, 500 mL of LB-Kan was inoculated with 1/100 of the overnight culture. Expression was induced at OD_600_ of 1.0 with 1 mM isopropyl β-D-1-thiogalactopyranoside (IPTG) for 12-14 h at 30 °C with vigorous agitation. Then, cells were harvested by centrifugation at 5,000 × g for 15 min at 4 °C, and the cellular pellet was resuspended in 50 mM Tris pH 8.0, 20 mM imidazole. The cell suspension was subjected to seven pulses of sonication (20 kc, 1 min) in an ice bath. The water-soluble fraction of this homogenate was recovered by centrifugation at 17,000 × g for 30 min at 4 °C. Then, 1 mL of HisPur Ni-NTA Resin (Thermo Fisher) equilibrated in resuspension buffer was added to the soluble fraction. The resin interacted with the water-soluble fraction end-over-end rotation overnight at 4 °C. The non-retained fraction was collected, and the resin was washed with 50 mM Tris pH 8.0, 20 mM imidazole until absorbance at 280 nm was below 0.05. Finally, the recombinant toxins were eluted with 5 column volumes of 50 mM Tris pH 8.0, 250 mM imidazole. Milligram amounts of protein samples were purified to homogeneity according to their electrophoretic behaviour analysed by SDS-PAGE. The recombinant toxins were dialyzed against PBS with 10,000 MWCO dialysis tubing (Snakeskin. Thermo Fisher), aliquoted, and flash-frozen with N_2_(l) for long-term storage at -20 °C.

### Reversed-phase high-performance liquid chromatography

The *H. lepturus* venom used in this study was provided by Seyed Mahdi Kazemi from Zagros Herpetological Institute and Mahboubeh Sadat Hosseinzadeh from the Department of Biology, Faculty of Sciences, University of Birjand, Birjand, Iran. To purify the native SMase from the venom, reversed-phase high-performance liquid chromatography (RP-HPLC) was performed using an Agilent Infinity II (Santa Clara, CA, USA) system, as previously described ^30^. Briefly, lyophilized venom (10 mg) was dissolved in 1 mL of water containing 0.1% trifluoroacetic acid (TFA; solution A), centrifuged at 14,000 × g for 10 min, and transferred to an HPLC vial. For each fractionation round, 100 µL of sample was injected into an RP-HPLC C_18_ column (250 × 4.6 mm, 5 µm particle size) and eluted at 1 mL/min using a gradient towards acetonitrile containing 0.1% TFA (solution B) (0–15% B for 15 min, 15–45% B for 60 min, 45–70% B for 10 min, and 70% B for 9 min). Fractions with an absorbance response (280 nm) 10 times above the background were analyzed by SDS-PAGE to detect the SMase based on its electrophoretic mobility. The fraction with the correct size was dialyzed against phosphate-buffered saline (PBS: 137 mM NaCl, 3 mM KCl, 8 mM Na_2_HPO_4_.2H_2_O, 1.4 mM KH_2_PO_4_, pH 7.4) with 10,000 MWCO dialysis tubing (Snakeskin, Thermo Fisher) and flashfrozen with N_2_(l) for long term storage at -20 °C.

### Near-UV circular dichroism

Circular dichroism experiments were performed using a Jasco J-810 circular dichroism spectropolarimeter, equipped with a Peltier-controlled cuvette holder. Samples of recombinant proteins (rLr, rHl, and cSMase) at a concentration of 0.3 mg/mL in PBS were loaded into quartz cuvettes with a sample length of 1 mm prior to measurement.

### Nanotemper

Protein denaturation analysis was conducted using a Nanotemper Prometheus NT.48 instrument, which employs microscale thermophoresis (MST) technology to monitor changes in protein fluorescence in response to temperature. Samples of proteins (1 μM in PBS) were heated from 20 °C to 95 °C at a rate of 2 °C/min, with changes in intrinsic protein fluorescence being measured using a NanoTemper capillary. Fluorescence data was collected at 1 °C intervals, and the melting temperature (T_m_) was calculated using the Prometheus NT.48 software, which fits the data to a Boltzmann sigmoidal curve, and calculated based on four independent, averaged experiments.

### Toxin biotinylation

The toxins were biotinylated using a 10:1 molar ratio of biotinylation reagent (Innolink Biotin 354S, Merck) to toxin as recommended by the manufacturer. Briefly, the biotinylation reagent was dissolved in dimethyl sulfoxide (DMSO) and added to the recombinant toxins in a small volume (< 5% of the final volume). After 90 min of incubation at 25 °C, biotinylated toxins were purified using Amicon® Ultra-4 Centrifugal Filter Units with a 10 kDa membrane, with four washes using 4 mL PBS at 8 °C. The protein concentration was determined by measuring absorbance at 280 nm with a NanoDrop and adjusted using the theoretical extinction coefficient calculated based on the protein sequence with ProtParam (Expasy)^31^. The degree of biotinylation was analyzed by the A280/A354 absorbance ratio, resulting in a 2:1 to 3:1 biotin:toxin molar ratio for all the toxins tested.

### Phage display selection

The IONTAS library, a human antibody phage display library of 4 × 10^10^ clones, was used for phage display selection. The library is constructed from B lymphocytes collected from 43 non-immunized human donors, and contains antibodies in the form of scFvs ^32^. Before each round of selection, streptavidin-specific phages were deselected using streptavidin-coated Dynabeads (Invitrogen, M-280). The biotinylated recombinant toxins were then incubated with the phage library in solution, which was blocked with 3% (w/v) skimmed milk in PBS (3MPBS). The biotinylated recombinant toxins were captured using streptavidin-coated Dynabeads (Invitrogen, M-280). After washing with PBS containing 0.1% (v/v) Tween 20 in a King Fisher Flex System, phages were eluted with trypsin and were added to cultures of *E. coli* TG1 cells with an OD_600_ of 0.5 and shaken at 150 rpm at 37 °C for 1 h. Then, the cultures were plated and incubated overnight at 30 °C on 2xTY medium plates (16 g/L Tryptone, 10 g/L yeast extract, 5 g/L NaCl, 15 g/L agar) supplemented with 2% glucose and 100 µg/mL ampicillin. The following day, the number of colony-forming units was determined and the plates were scraped using 2 mL of 2xTY medium (16 g/L Tryptone, 10 g/L yeast extract, 5 g/L NaCl), supplemented with 2% (w/v) glucose and 50 μg/mL kanamycin and 25 % (v/v) glycerol. The aliquoted phages were stored at -80 °C.

Two panning strategies were compared: 1) Stringent panning with cSMase, in which three rounds of panning were performed, while the antigen concentration was reduced in every round (100, 50, and 10 nM), and 2) cross-panning with recombinant SMases from *H. lepturus* (rHl) and *L. rufescens* (rLr), in which the antigen was switched in the second and third round of panning, with the same antigen concentration of 100 nM in round 2, but with a lower antigen concentration of 50 nM in round 3 (i.e., rLr 100 nM, rHl 100 nM, rHl 50 nM and rHl 100 nM, rLr 100 nM, rLr 50 nM). To recover phages from the selection outputs, 5 mL of 2xTY medium plates (16 g/L Tryptone, 10 g/L yeast extract, 5 g/L NaCl) with 2% glucose and 100 µg/mL ampicillin were inoculated with the glycerol stock and grown at 37 °C under shaking at 280 rpm until the OD_600_ reached 0.5. Then, the cultures were inoculated with 20x coverage M13KO7 helper phage ^33^, which confers resistance to kanamycin, and incubated for 1 hour at 37 °C with shaking at 150 rpm. Then, the cultures were centrifuged at 3,500 × g for 10 minutes at 4 °C, and the pellet was resuspended in 2xTY medium supplemented with 100 µg/mL ampicillin and 50 µg/mL kanamycin. The cultures were shaken overnight at 25 °C at 280 rpm to produce phages.Rescued phages from every round were precipitated with 20% (w/v) PEG-8000, 2.5 M NaCl, followed by centrifugation and subsequent resuspension in PBS. Rescued phages were kept at 4 °C for less than a week. 15% (v/v) glycerol was added to the phage aliquots for long term storage at -80 °C.

### Polyclonal phage ELISA

The selected phage outputs were evaluated for antigen binding using a polyclonal phage ELISA similarly to what has previously been described ^32^. In short, clear Maxisorp plates (Nunc) were coated with streptavidin (10 mg/mL) and biotinylated antigens (150 nM) were indirectly immobilized on the streptavidin-coated wells. Negative control wells were coated with streptavidin or an irrelevant biotinylated toxin (α-cobratoxin from *Naja kaouthia*), while positive control wells were coated with M13 helper phages in PBS. The plates were washed three times with PBS containing 0.1% (v/v) Tween 20 and then three times with PBS, before the addition of phage outputs from the different rounds in 3MPBS. After one hour of incubation, plates were washed as before and 1:2,000 anti-M13-HRP antibody (Sino Biological #11973-MM05T-H) in 3MPBS was added to the wells. Following one hour of incubation and a final washing step, the plates were incubated with 3,3’,5,5’-tetramethylbenzidine (TMB) (Thermo Fisher #34021) for 15-20 min, followed by the addition of 0.5 M H_2_SO_4_ to stop the reaction. Binding signals were detected by measuring the absorbance at 450 nm.

### Subcloning

The scFv genes from the third selection round were amplified using PCR and subcloned using *NcoI* and *NotI* restriction endonuclease sites into the pSANG10-3F vector for expression of soluble scFvs transformed into *E. coli* BL21(DE3) cells (New England Biolabs), as previously descrived [6,10]. The pSANG10-3F vector contains a FLAG-tag and a 6xHis tag in phase with the scFv. For each of the third selection rounds from the stringent consensus phage display campaign, 276 individual scFv clones were picked and incubated at 30 °C and 800 rpm in 96-well plates overnight in 1 mL of autoinduction medium. The next day, the scFv-containing supernatants were tested for binding in an expression-normalized capture DELFIA, as previously described ^32^.

### scFv expression and purification

The TPL0674_03_F05 and TPL0674_03_A04 scFv clones were produced and purified as previously described ^5^. Briefly, 2xTY medium, supplemented with 2% (w/v) glucose and 50 μg/mL kanamycin was inoculated with TPL0674_03_F05 and TPL0674_03_A04. The cultures were incubated overnight at 37 °C and 250 rpm. The following day, 500 mL of autoinduction medium was inoculated with the overnight cultures and further incubated overnight at 30 °C and 200 rpm. The cells were then harvested by centrifugation at 4,300 × g for 10 min, and the supernatants were discarded. The cell pellet was resuspended in 50 mL of TES-buffer (30 mM Tris–HCl pH 8.0, 1 mM EDTA, 20% sucrose (w/v)) containing 1.5 kU/mL of r-lysozyme (∼ 70,000 U/mg). After 20 min of incubation on ice, the cells were centrifuged at 4,300 × g for 10 min, and the supernatant was discarded. The cell pellet was resuspended in 50 mL of 5 mM MgSO_4_ supplemented with the same amount of r-lysozyme as above and incubated on ice for 20 min. After centrifugation at 4,300 × g for 10 minutes, the supernatant was pooled with the supernatant from the previous step and kept on ice. The pooled supernatants were then centrifuged at 30,000 × g for 30 min. The His-tagged scFvs were purified using the same method as above for the recombinant toxins.

### Expression-normalized capture DELFIA

The binding of the scFvs to biotinylated rHl and rLr (30 nM) was assessed using a DELFIA-based assay on black Maxisorp plates (Nunc), as previously described ^32^. Briefly, the scFvs were immobilised with anti-FLAG M2 antibody (Sigma-Aldrich #F3165) at 2.5 µg/mL. After immobilization of the scFvs, the biotinylated toxins (rHl and rLr) were added in 3MPBS. Binding was detected using streptavidin conjugated with europium (Perkin Elmer #1244-360) and DELFIA Enhancement Solution (Perkin Elmer, 4001–0010).

### Biolayer interferometry

Biolayer interferometry analysis was performed using a ForteBio Octet RED96 instrument with streptavidin tips. The tips were hydrated in kinetics buffer (Sartorius) for 30 min prior to use, then washed with PBS to remove any non-specifically bound proteins. Biotinylated, recombinant SMase from *L. rusfescens* (rLr), recombinant SMase from *H. lepturus* (rHl), or natural SMase purified from *H. lepturus* venom (Hl) were immobilized onto the tips in kinetics buffer at 50 nM for 10 min. Binding experiments were performed by dipping the streptavidin-coated tips into a 96-well microplate containing the purified scFvs at 100, 250, and 500 nM, followed by a dissociation step in kinetics buffer. Data analysis was performed using ForteBio Data Analysis software. The association and dissociation rate constants (k_a_ and k_d_) were calculated using a 1:1 binding model. The affinity constant (K_D_) was calculated as the ratio of k_d_ to k_a_. All data was analyzed in triplicate, and the average values were reported.

### SMase activity neutralization assay

The assess the ability of the TPL0674_03_F05 and TPL0674_03_A04 scFvs clones to neutralize *H. lepturus* whole venom, a Colorimetric Sphingomyelinase Assay Kit (#MAK152-1KT, ThermoFisher) was used according to the manufacturer’s specifications. The experiment was independently performed twice, with duplicates for each well. Briefly, the scFvs were diluted to concentrations ranging from 0.5-8 μM and then incubated with *H. lepturus* venom at a final concentration of 1.5 mg/mL. The mixture was combined with an assay reaction mix containing sphingomyelin and incubated for 2 h at 37 °C. An SMase assay mixture containing Alkaline phosphatase (ALP) and other enzymes was added to each well. The dephosphorylation of phosphorylcholine to choline by ALP results in the production of a colored product. The SMase activity was quantified by detecting the endpoint absorbance at 655 nm. The assay was run in parallel with a calibration curve, and negative controls included buffer only and a non-binding scFv (TPL0684_01_C09). The amount of sphingomyelinase activity present in the samples was determined from the standard curve.

## Acknowledgements

The authors are supported by a grant from the European Research Council (ERC) under the European Union’s Horizon 2020 research and innovation programme [grant no. 850974]. Thank you to Shirin Ahmadi for providing the contact with SMK to obtain the *H. lepturus* venom.

## Author contribution

ERdT, MB, and AHL conceptualized the project, ERdT designed the experiments, ERdT, and SL performed the experiments, ERdT, MB, and AHL, drafted and redacted the manuscript, SMK extracted and provided the *H. lepturus* venom, all the authors contributed to write, the manuscript.

## Abbreviations

bnAbs: Broadly-neutralising antibodies
cSMase: Consensus Sphingomyelinase
Hl: *Hemiscorpius lepturus* venom-derived Sphingomyelinase
mAbs: Monoclonal antibodies
rHl: Recombinant *Hemiscorpius lepturus* Sphingomyelinase
rLr: Recombinant *Loxosceles rufescens* Sphingomyelinase
scFv: Single-chain variable fragment
SMase: Sphingomyelinase

## Supplementary material

**File S1.** List of the 136 protein sequences used to design the cSMase sequence.

## References

1. Kaplon H, Chenoweth A, Crescioli S, Reichert JM (2022) Antibodies to watch in 2022. mAbs 14:2014296.

2. Salazar G, Zhang N, Fu T-M, An Z (2017) Antibody therapies for the prevention and treatment of viral infections. Npj Vaccines 2:1–12.

3. Wilhelm A, Widera M, Grikscheit K, Toptan T, Schenk B, Pallas C, Metzler M, Kohmer N, Hoehl S, Marschalek R, et al. (2022) Limited neutralisation of the SARS-CoV-2 Omicron subvariants BA.1 and BA.2 by convalescent and vaccine serum and monoclonal antibodies. eBioMedicine 82:104158.

4. Casewell NR, Jackson TNW, Laustsen AH, Sunagar K (2020) Causes and Consequences of Snake Venom Variation. Trends Pharmacol. Sci. 41:570–581.

5. Ahmadi S, Pucca M, Jürgensen JA, Janke R, Schoof E, Sørensen C, Caliskan F, Laustsen A (2020) An in vitro methodology for discovering broadly-neutralizing monoclonal antibodies. Sci. Rep. 10.

6. Ledsgaard L, Wade J, Jenkins TP, Boddum K, Oganesyan I, Harrison JA, Villar P, Leah RA, Zenobi R, Schoffelen S, et al. (2023) Discovery and optimization of a broadly-neutralizing human monoclonal antibody against long-chain α-neurotoxins from snakes. Nat. Commun. 14:682.

7. Rodríguez-Rodríguez ER, Olamendi-Portugal T, Serrano-Posada H, Arredondo-López JN, Gómez-Ramírez I, Fernández-Taboada G, Possani LD, Anguiano-Vega GA, Riaño-Umbarila L, Becerril B (2016) Broadening the neutralizing capacity of a family of antibody fragments against different toxins from Mexican scorpions. Toxicon 119:52–63.

8. Glanville J, Andrade JC, Bellin M, Kim S, Pletnev S, Tsao D, Verardi R, Bedi R, Friede T, Liao S, et al. (2022) Venom protection by antibody from a snakebite hyperimmune subject. :2022.09.26.507364. Available from: https://www.biorxiv.org/content/10.1101/2022.09.26.507364v1

9. Rivera-de-Torre E, Rimbault C, Jenkins TP, Sørensen CV, Damsbo A, Saez NJ, Duhoo Y, Hackney CM, Ellgaard L, Laustsen AH (2022) Strategies for Heterologous Expression, Synthesis, and Purification of Animal Venom Toxins. Front. Bioeng. Biotechnol. [Internet] 9. Available from: https://www.frontiersin.org/article/10.3389/fbioe.2021.811905

10. de la Rosa G, Olvera F, Archundia IG, Lomonte B, Alagón A, Corzo G (2019) Horse immunization with short-chain consensus α-neurotoxin generates antibodies against broad spectrum of elapid venomous species. Nat. Commun. 10:3642.

11. de la Rosa G, Corrales-García LL, Rodriguez-Ruiz X, López-Vera E, Corzo G (2018) Short-chain consensus alpha-neurotoxin: a synthetic 60-mer peptide with generic traits and enhanced immunogenic properties. Amino Acids 50:885–895.

12. Laustsen AH, Greiff V, Karatt-Vellatt A, Muyldermans S, Jenkins TP (2021) Animal Immunization, in Vitro Display Technologies, and Machine Learning for Antibody Discovery. Trends Biotechnol. 39:1263–1273.

13. Ledsgaard L, Ljungars A, Rimbault C, Sørensen CV, Tulika T, Wade J, Wouters Y, McCafferty J, Laustsen AH (2022) Advances in antibody phage display technology. Drug Discov. Today 27:2151–2169.

14. Jenkins TP, Ahmadi S, Bittenbinder MA, Stewart TK, Akgun DE, Hale M, Nasrabadi NN, Wolff DS, Vonk FJ, Kool J, et al. (2021) Terrestrial venomous animals, the envenomings they cause, and treatment perspectives in the Middle East and North Africa. PLoS Negl. Trop. Dis. 15:e0009880.

15. Dehghani R, Fathi B (2012) Scorpion sting in Iran: A review. Toxicon 60:919–933.

16. de Oliveira KC, Gonçalves de Andrade RM, Piazza RMF, Ferreira JMC, van den Berg CW, Tambourgi DV (2005) Variations in Loxosceles spider venom composition and toxicity contribute to the severity of envenomation. Toxicon 45:421–429.

17. Catalán A, Cortes W, Sagua H, González J, Araya JE (2011) Two new phospholipase D isoforms of Loxosceles laeta: Cloning, heterologous expression, functional characterization, and potential biotechnological application. J. Biochem. Mol. Toxicol. 25:393–403.

18. Seyedian R, Pipelzadeh MH, Jalali A, Kim E, Lee H, Kang C, Cha M, Sohn E, Jung E-S, Rahmani AH, et al. (2010) Enzymatic analysis of Hemiscorpius lepturus scorpion venom using zymography and venom-specific antivenin. Toxicon 56:521–525.

19. de Andrade SA, Pedrosa MFF, de Andrade RMG, Oliva MLV, van den Berg CW, Tambourgi DV (2005) Conformational changes of Loxosceles venom sphingomyelinases monitored by circular dichroism. Biochem. Biophys. Res. Commun. 327:117–123.

20. Martin CD, Rojas G, Mitchell JN, Vincent KJ, Wu J, McCafferty J, Schofield DJ (2006) A simple vector system to improve performance and utilisation of recombinant antibodies. BMC Biotechnol. 6:46.

21. Laustsen AH Antivenom in the Age of Recombinant DNA Technology. In: Handbook of Venoms and Toxins of Reptiles. 2nd ed. CRC Press; 2021.

22. Campbell BB, Light N, Fabrizio D, Zatzman M, Fuligni F, de Borja R, Davidson S, Edwards M, Elvin JA, Hodel KP, et al. (2017) Comprehensive Analysis of Hypermutation in Human Cancer. Cell 171:1042–1056.e10.

23. Wendel BS, Fu Y, He C, Hernandez SM, Qu M, Zhang Z, Jiang Y, Han X, Xu J, Ding H, et al. (2020) Rapid HIV Progression Is Associated with Extensive Ongoing Somatic Hypermutation. J. Immunol. Author Choice 205:587–594.

24. Woodford N, Ellington MJ (2007) The emergence of antibiotic resistance by mutation. Clin. Microbiol. Infect. 13:5–18.

25. Waterhouse AM, Procter JB, Martin DMA, Clamp M, Barton GJ (2009) Jalview Version 2— a multiple sequence alignment editor and analysis workbench. Bioinformatics 25:1189–1191.

26. Jumper J, Evans R, Pritzel A, Green T, Figurnov M, Ronneberger O, Tunyasuvunakool K, Bates R, Žídek A, Potapenko A, et al. (2021) Highly accurate protein structure prediction with AlphaFold. Nature 596:583–589.

27. Mirdita M, Schütze K, Moriwaki Y, Heo L, Ovchinnikov S, Steinegger M (2022) ColabFold: making protein folding accessible to all. Nat. Methods 19:679–682.

28. Varadi M, Anyango S, Deshpande M, Nair S, Natassia C, Yordanova G, Yuan D, Stroe O, Wood G, Laydon A, et al. (2022) AlphaFold Protein Structure Database: massively expanding the structural coverage of protein-sequence space with high-accuracy models. Nucleic Acids Res. 50:D439–D444.

29. Liebschner D, Afonine PV, Baker ML, Bunkóczi G, Chen VB, Croll TI, Hintze B, Hung L- W, Jain S, McCoy AJ, et al. (2019) Macromolecular structure determination using X-rays, neutrons and electrons: recent developments in Phenix. Acta Crystallogr. Sect. Struct. Biol. 75:861–877.

30. Calvete JJ (2011) Proteomic tools against the neglected pathology of snake bite envenoming. Expert Rev. Proteomics 8:739–758.

31. Walker JM ed The Proteomics Protocols Handbook. Totowa, NJ: Humana Press; 2005. Available from: http://link.springer.com/10.1385/1592598900

32. Laustsen AH, Karatt-Vellatt A, Masters EW, Arias AS, Pus U, Knudsen C, Oscoz S, Slavny P, Griffiths DT, Luther AM, et al. (2018) In vivo neutralization of dendrotoxin-mediated neurotoxicity of black mamba venom by oligoclonal human IgG antibodies. Nat. Commun. 9:1–9.

33. Jensen L, Kilstrup M, Karatt-Vellatt A, Mccafferty J, Laustsen A (2018) Basics of Antibody Phage Display Technology. Toxins 10:236.

